# MYC and AP-1 oncogenes synergistically bind enhancers to rewire transcription

**DOI:** 10.1101/2025.04.28.650480

**Authors:** Reshma Kalyan Sundaram, Ravi Radhakrishnan, Bomyi Lim

## Abstract

The transcription factor c-MYC (MYC) is deregulated in about 70% of human cancers. Through *de novo* motif discovery analysis on published MYC ChIP-seq datasets from cancer cell lines, we found cell-type specific co-enrichment of the TRE motifs (AP-1 binding sites), alongside MYC’s canonical EBOX motif. TRE motifs serve as indirect MYC binding sites in synergy with AP-1, predominantly at enhancers rather than promoters. At elevated MYC levels, as seen in cancers, MYC preferentially occupies TRE sites over EBOX sites at enhancers. Integration of ChIP-seq and RNA-seq data revealed TRE enhancer binding sites are frequently associated with MYC-mediated transcriptional repression. Gene Ontology analysis showed that MYC utilizes TRE sites to transcriptionally rewire cells, modulating cancer hallmarks like proliferation, apoptosis, and cell adhesion. These molecular insights into how increased MYC levels alter its DNA binding preference and gene regulation could inform new therapeutic strategies targeting cancer-specific MYC functions and its co-regulators.

## Introduction

The *MYC* family of proto-oncogenes comprises of *c-MYC, N-MYC* and *L-MYC* (Dang, 2012; Dhanasekaran et al., 2022; Jakobsen & Siersbæk, 2025). These genes code for the transcription factors c-MYC (MYC), N-MYC (MYCN) and L-MYC (MYCL) respectively (Dhanasekaran et al., 2022; Jakobsen & Siersbæk, 2025; Kress et al., 2015). Here, we focus on transcription factor (TF) MYC, which was identified over 40 years ago (Colby et al., 1983; Meyer & Penn, 2008; Vennstrom et al., 1982). MYC is expressed in various cell and tissue types, and drives both general as well as cell-type specific functions (Dang, 2012; Dhanasekaran et al., 2022; Jakobsen & Siersbæk, 2025). The genes regulated by MYC correspond to several important cellular processes such as cell cycle, cell growth, metabolism, DNA replication and apoptosis (Kress et al., 2015). Additionally, MYC-regulated genes and pathways are known to indirectly regulate the global transcriptome through feedback mechanisms that modulate RNA synthesis (Kress et al., 2015). Since MYC plays such a pivotal role as a master regulator of the transcriptome, MYC levels in cells are tightly controlled to ensure normal cellular function (Dang, 2012; Meyer & Penn, 2008).

MYC, however, is deregulated in up to 70% of human cancers (Dang, 2012) through mechanisms such as gene amplification, signal transduction, and protein stabilization (Dhanasekaran et al., 2022). These processes elevate MYC levels in cells, triggering the activation of new transcriptional programs by MYC and causing cellular transformation (Kress et al., 2015). Such essential roles of MYC in cancer initiation and maintenance (Dhanasekaran et al., 2022; Gabay et al., 2014) establish it as an important target of interest in cancer treatments (Dhanasekaran et al., 2022; Llombart & Mansour, 2022; Lourenco et al., 2021). Several approaches to target MYC have been explored, including silencing MYC gene expression, inhibiting MYC translation, promoting MYC protein degradation, and disrupting MYC protein-protein interactions (Dhanasekaran et al., 2022; Llombart & Mansour, 2022; Lourenco et al., 2021; Whitfield & Soucek, 2025; Wolf & Eilers, 2020). However, no MYC inhibitors are currently available for clinical use (Whitfield & Soucek, 2025).

In cancers, MYC’s TF function is altered through changes in its binding patterns (Jakobsen & Siersbæk, 2025; Kress et al., 2015; Lorenzin et al., 2016), regulated gene targets (Jakobsen & Siersbæk, 2025; Kress et al., 2016; Lorenzin et al., 2016; Sabò et al., 2014; Walz et al., 2014), and interactions with co-regulator proteins (Jakobsen & Siersbæk, 2025; Lourenco et al., 2021; Tu et al., 2015). Chromatin immunoprecipitation followed by sequencing (ChIP-seq) technology (Barski et al., 2007; Johnson et al., 2007; Mikkelsen et al., 2007; Robertson et al., 2007) enables genome-wide analysis of MYC’s DNA binding patterns. Additionally, integrating ChIP-seq with RNA sequencing (RNA-seq) (Pellanda et al., 2021; Walz et al., 2014) allows for the identification of MYC-regulated target genes. ChIP-seq can also be used to infer regulatory complex formation through MYC’s interactions with other proteins (Dunham et al., 2012; Feldker et al., 2020), providing insights into its broader regulatory network.

Therefore, in this work, we analyzed publicly-available ChIP-seq datasets (Dunham et al., 2012; Hitz et al., 2023; Kagda et al., 2023; Luo et al., 2019; Muthalagu et al., 2014; Walz et al., 2014) of various cancer cell lines to gain a comprehensive understanding of MYC’s oncogenic gene regulatory functions. Our analysis revealed a surprising co-enrichment of the TRE motif, a known AP-1 TF family binding site (Eferl & Wagner, 2003; Garces de los Fayos Alonso et al., 2018), alongside the canonical EBOX motif (Jakobsen & Siersbæk, 2025; Kress et al., 2015). Subsequent analysis of MYC and AP-1 ChIP-seq datasets (Dunham et al., 2012; Hitz et al., 2023; Kagda et al., 2023; Luo et al., 2019) showed that the TRE motif serves as an indirect MYC binding site, occupied in synergy with AP-1 TFs. Additionally, we found that the TRE motif is an enhancer-specific MYC binding site, unlike the EBOX motif which is both an enhancer and promoter MYC binding site. Analysis of MYC ChIP-seq datasets across different MYC expression levels (Muthalagu et al., 2014; Walz et al., 2014) confirmed that TRE remains an enhancer-specific MYC binding site independent of MYC levels in cells. Gene Ontology (Huang da et al., 2009; Sherman et al., 2022) on MYC target genes revealed that MYC utilizes TRE binding sites at enhancers to transcriptionally rewire cells, by modulating several cancer hallmarks (Dhanasekaran et al., 2022; Hanahan & Weinberg, 2000; Hanahan & Weinberg, 2011) such as proliferation, apoptosis, and cell adhesion. Our findings will potentially aid in development of new treatments targeting cancer-specific functions of MYC and its co-regulators.

## Results

### Genome-wide analysis reveals that MYC indirectly occupies TRE binding site in synergy with AP-1

Genome-wide DNA binding patterns of MYC were identified through *de novo* motif discovery (Heinz et al., 2010) using ENCODE (Dunham et al., 2012; Hitz et al., 2023; Kagda et al., 2023; Luo et al., 2019) MYC ChIP-seq datasets from 11 cell lines. The EBOX motif (CACGTG) emerged as the first ranked significantly enriched motif in the MYC ChIP-seq datasets across all analyzed cell lines (Fig. 1a,b), consistent with its well-established role as a direct consensus binding site for MYC (Jakobsen & Siersbæk, 2025; Kress et al., 2015). Surprisingly, the TRE motif (TGA(G/C)TCA) also appeared among the top 6 ranked significantly enriched motifs in the MYC ChIP-seq datasets of 8 out of the 11 analyzed cell lines (Fig. 1a,b). The TRE motif is a well-known binding site for the AP-1 family of TFs (Eferl & Wagner, 2003; Garces de los Fayos Alonso et al., 2018).

**Fig. 1:**
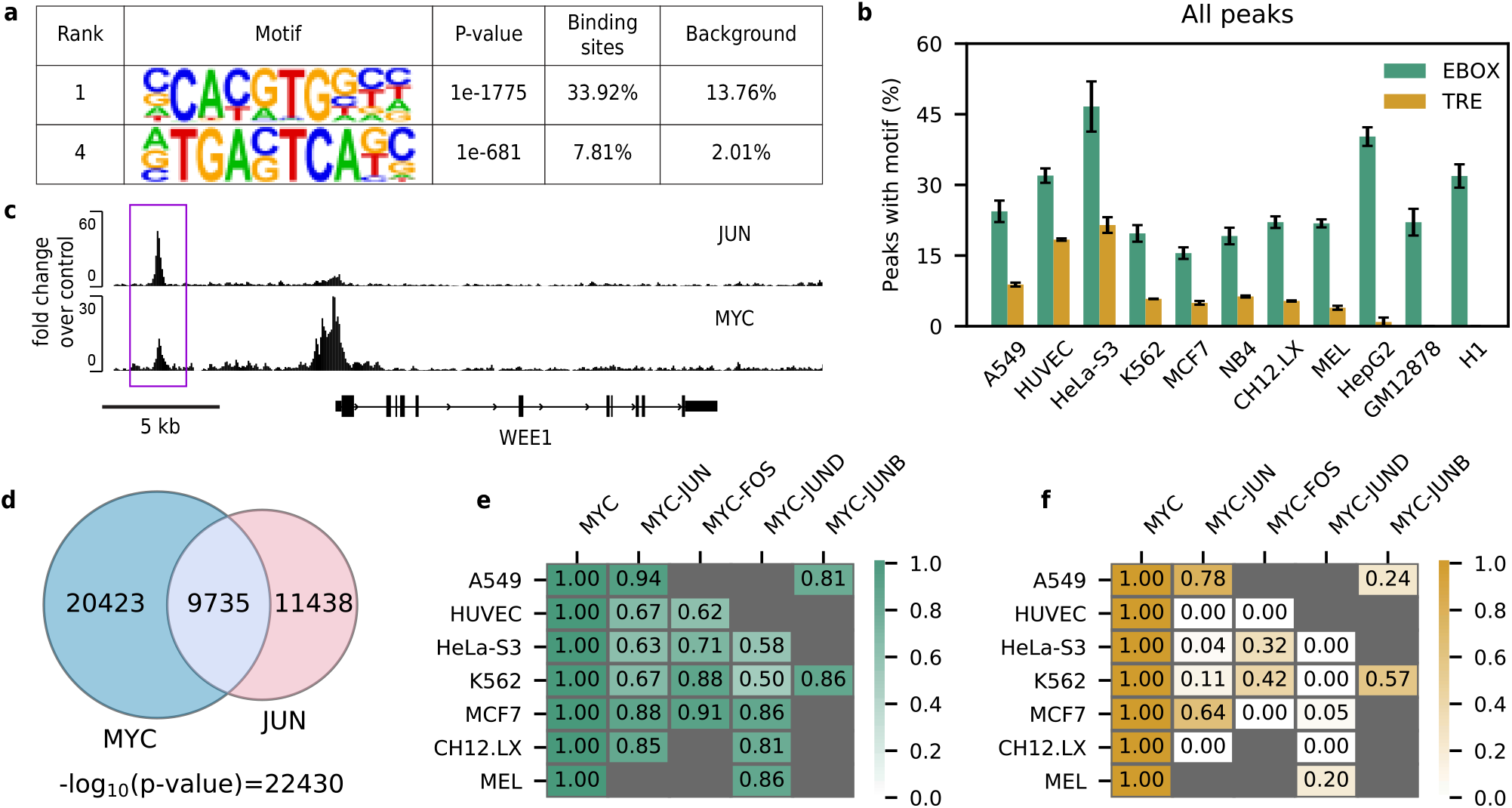
Uncovering MYC and AP-1 synergy in DNA-binding in various cancer cell lines. **a.** Enrichment of EBOX and TRE motifs in MYC ChIP-seq dataset of a representative cell line, K562, using the Homer *de novo* motif enrichment analysis. P-value calculated using binomial enrichment test. **b**. Quantification of EBOX and TRE motif enrichment in MYC ChIP-seq datasets for various cell lines analyzed. Data represents mean and standard deviation of 4 replicates. **c**. An example of overlapping MYC and an AP-1 family member JUN ChIP-seq peaks near WEE1 gene for a representative cell line, K562. **d**. Overlapping DNA binding regions of MYC and JUN, an AP-1 family member, identified using ChIP-seq datasets for a representative cell line, K562. P-value of overlap calculated using a hypergeometric test. Proportion of MYC ChIP-seq peaks containing **e**. EBOX and **f**. TRE motifs (normalized per cell line) before and after the removal of co-bound peaks with various AP-1 family TFs in multiple cell lines. The drastic reduction in MYC ChIP-seq peaks containing TRE motifs upon removal of peaks overlapping with AP-1 ChIP-seq shows TRE motifs enriched in MYC ChIP-seq as a synergistic binding site of MYC with AP-1 TFs.

Since ChIP-seq captures genome-wide binding patterns of a TF of interest (Barski et al., 2007; Johnson et al., 2007; Mikkelsen et al., 2007; Robertson et al., 2007), including both direct and indirect binding regions (Liang et al., 2014; Neph et al., 2012; Ronzio et al., 2020; Wang et al., 2012), *de novo* motif discovery can reveal co-enriched motifs alongside the consensus motif, as observed in previous studies (Feldker et al., 2020; Heinz et al., 2010; Ronzio et al., 2020; Wang et al., 2012). Co-enriched motifs include DNA regions indirectly bound by the TF, typically due to the formation of regulatory complexes or synergistic interactions with other proteins (Feldker et al., 2020; Heinz et al., 2010; Ronzio et al., 2020; Wang et al., 2012). Given that the 1) TRE motif is a known consensus binding site for AP-1 (Eferl & Wagner, 2003; Garces de los Fayos Alonso et al., 2018), 2) TRE motif sequence significantly differs from MYC’s consensus binding motif (EBOX) sequence, and 3) TRE enrichment was consistently lower than EBOX enrichment in the MYC ChIP-seq datasets across all analyzed cell lines (Fig. 1b), we hypothesized that TRE could represent an indirect binding site for MYC, occupied in synergy with AP-1.

To test this hypothesis, we first examined whether MYC and AP-1 family TFs shared overlapping genomic binding regions. In K562 cells, ∼30% of MYC ChIP-seq peaks overlapped with peaks of the AP-1 family member JUN, which itself showed ∼50% overlap with MYC peaks (Fig 1c, d). We subsequently analyzed the overlap of ChIP-seq datasets for MYC and 4 AP-1 family TFs (JUN, FOS, JUND, and JUNB) across 7 cell lines (Fig. 1d, Extended Data Fig. 1a) (Dunham et al., 2012; Hitz et al., 2023; Kagda et al., 2023; Luo et al., 2019). Remarkably, we observed statistically significant overlap between ChIP-seq datasets for all combinations of MYC and AP-1 TFs across all analyzed cell lines (Fig. 1d, Extended Data Fig. 1a). These findings confirmed that MYC and AP-1 indeed bind to overlapping genomic locations.

We then examined if the observed overlap between MYC and AP-1 TFs binding may explain the enrichment of the TRE motif in the MYC ChIP-seq datasets. We quantified the number of EBOX and TRE containing MYC ChIP-seq peaks (referred to as “occurrence”) that do not overlap with the AP-1 family member ChIP-seq peaks. Specifically, using ChIP-seq datasets for 4 AP-1 family TFs across 7 cell lines, we first calculated the occurrence of EBOX and TRE motifs in the MYC ChIP-seq dataset for each cell line. We then recalculated the occurrence after removing peaks that overlapped with the ChIP-seq peaks for individual AP-1 TFs in the same cell line.

We found that the occurrence of TRE motif decreased drastically when overlapping regions with AP-1 TFs were individually removed from the MYC ChIP-seq datasets (Fig. 1f, Extended Data Fig. 1c). In most cell lines, this value even dropped to zero, depending on the specific AP-1 TFs available for analysis. These findings suggest that the enrichment of TRE motifs in MYC ChIP-seq datasets is predominantly due to synergistic binding with AP-1 TFs, supporting the hypothesis that TRE motifs represent indirect binding sites for MYC mediated by interactions with AP-1.

The occurrence of the EBOX motif also decreased across all cell lines upon removal of the overlapping peaks with AP-1 TFs (Fig. 1e, Extended Data Fig. 1b). We assume that this reduction corresponds to EBOX sites bound by MYC in synergy with AP-1 TFs. However, the magnitude of reduction in EBOX occurrence was moderate at best (Fig. 1e,f, Extended Data Fig. 1b,c), suggesting that a significant number of EBOX-containing MYC ChIP-seq peaks do not show evidence of synergy with AP-1 TFs.

The reduction in both EBOX and TRE occurrences upon removal of AP-1 overlap from MYC ChIP-seq (Fig. 1e,f, Extended Data Fig. 1b,c) suggests that both EBOX and TRE binding sites are utilized in the synergistic interaction between MYC and AP-1 TFs. Based on this observation, we propose that when MYC and AP-1 interact, they directly bind DNA at their respective consensus binding sites, EBOX and TRE. Additionally, these findings indicate that while MYC often engages in synergistic interactions with AP-1, MYC could also bind EBOX motifs independently of AP-1 TFs in many genomic regions.

### Regulatory element analysis reveals that MYC binds to TRE sites specifically at enhancers

Previous studies (Jakobsen et al., 2024; Jakobsen & Siersbæk, 2025; Lin et al., 2012; Walz et al., 2014) have established that MYC regulates gene expression by binding to both promoters and enhancers. Given that MYC performs distinct gene regulatory functions at promoters and enhancers (Jakobsen et al., 2024; Jakobsen & Siersbæk, 2025; Lin et al., 2012), we examined the differences in MYC binding patterns across these regions. This analysis was conducted for 7 cell lines, for which MYC, H3K4me3, and H3K27ac ChIP-seq datasets were available in the ENCODE database (Dunham et al., 2012; Hitz et al., 2023; Kagda et al., 2023; Luo et al., 2019).

MYC-bound transcriptionally active promoters and MYC-bound active enhancer regions for 7 cell lines were identified using the H3K4me3 and H3K27ac ChIP-seq datasets respectively (see Supplemental Methods). As expected, we observed MYC binding at both promoter (marked by H3K4me3) and enhancer (marked by H3K27ac) regions at individual gene locus (Fig. 2a,b), as well as on the genome-wide level (Fig. 2c, Extended Data Fig. 2a-b). We examined the fraction of MYC ChIP-seq peaks corresponding to promoters and enhancers across the 7 cell lines (Fig. 2d). We found that a significant percentage of peaks were associated with both promoters and enhancers in all analyzed cell lines, highlighting the critical role of MYC binding at these regulatory elements for its function (Jakobsen et al., 2024; Jakobsen & Siersbæk, 2025; Lin et al., 2012).

**Fig. 2:**
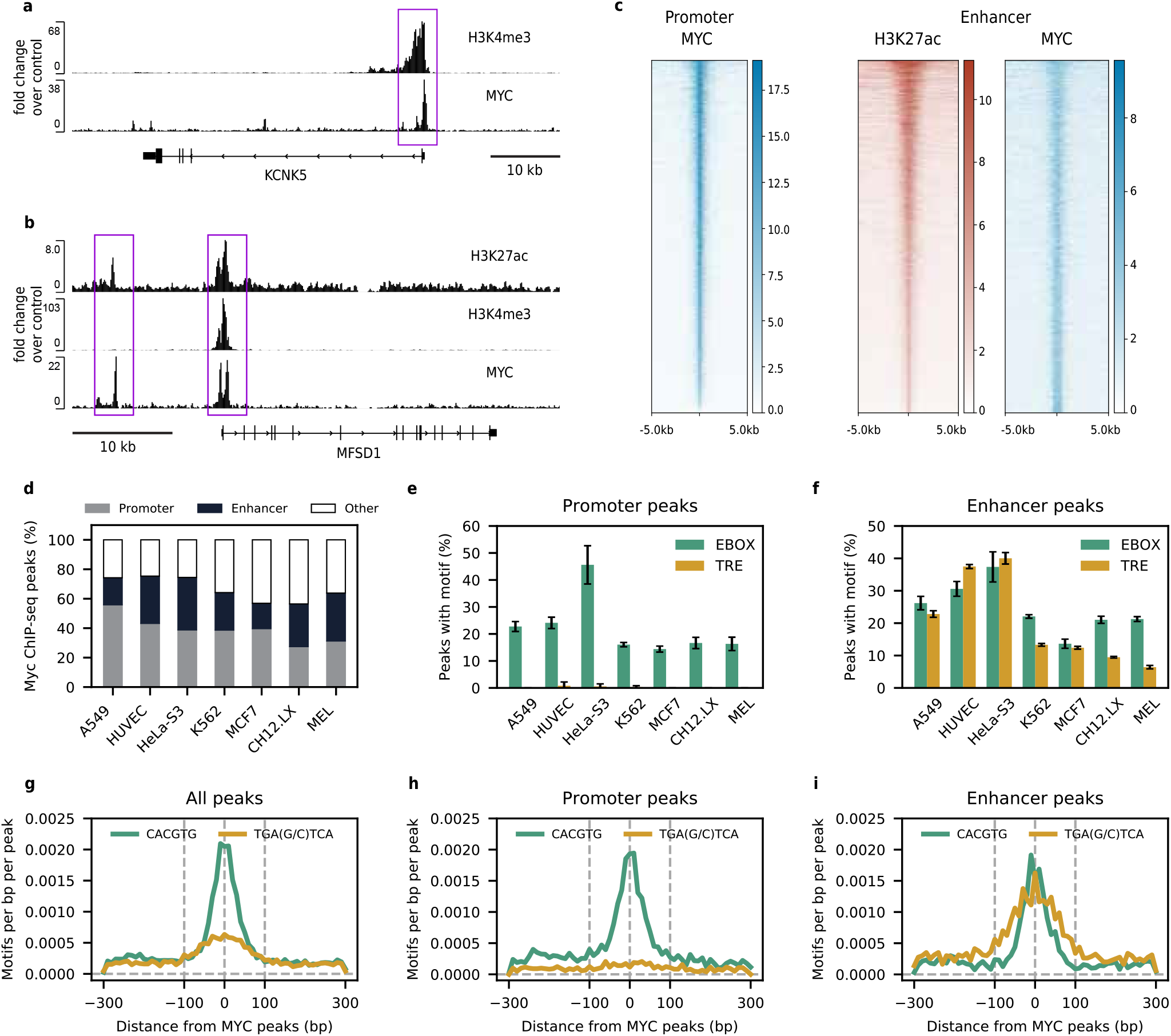
Identifying TRE as an enhancer-specific MYC binding site in various cancer cell lines. **a.** An example of overlapping MYC and H3K4me3 DNA binding regions identified from ChIP-seq near KCNK5 gene for K562 cell line. Boxed region with MYC and H3K4me3 peaks shows the identified MYC promoter binding site. **b**. An example of overlapping MYC and H3K27ac DNA binding regions identified from ChIP-seq near MFSD1 gene for K562 cell line. Boxed region on left with MYC and H3K27ac peaks shows the identified MYC enhancer binding site. Boxed region on right with MYC and H3K4me3 peaks shows the identified MYC promoter binding site. **c**. Heatmaps showing ChIP-seq binding intensities around promoter and enhancer regions of MYC for a representative cell line, K562. (Left) For promoters, MYC binding intensities are plotted around the center of TSS of actively transcribed genes bound by MYC. Rows are sorted in decreasing order of MYC binding intensities. (Right) For enhancers, MYC and H3K27ac binding intensities are plotted around the center of active enhancers bound by MYC. Rows are sorted in decreasing order of H3K27ac binding intensities. All heatmaps cover +/-5 kb window around the reference chosen for centering. Color bar indicates ChIP-seq binding intensities for corresponding heatmaps. **d**. Distribution of MYC binding to promoters, enhancers, and other genomic regions identified using MYC, H3K4me3, and H3K27ac ChIP-seq datasets of various cell lines. **e-f**. Enrichment of EBOX and TRE motifs in **e**. promoter and **f**. enhancer binding regions of MYC in various cell lines identified using the Homer *de novo* motif enrichment analysis. Data represents mean and standard deviation of 4 replicates. **g-i**. Density plot of EBOX and TRE motifs around the center of MYC binding regions identified from ChIP-seq experiment for K562 cell line. Dashed lines indicate +-100bp window around the center of MYC ChIP-seq peak used for performing *de novo* motif discovery analysis. Plots correspond to **g**. genome-wide, **h**. promoter, and **i**. enhancer regions of MYC binding.

We then performed *de novo* motif discovery on MYC ChIP-seq peaks at promoters and enhancers separately for the 7 cell lines (Fig. 2e,f). MYC’s consensus binding site, the EBOX motif, emerged as the first ranked significantly enriched motif in promoters, and among the top 4 ranked significantly enriched motifs in enhancers across all analyzed cell lines (Fig. 2e,f). However, the indirect MYC binding site identified in our study, the TRE motif, was among the top 5 ranked significantly enriched motifs only in enhancers but not in promoters across all analyzed cell lines (Fig. 2e,f). These findings suggest that while EBOX serves both as a promoter and enhancer binding site for MYC, TRE serves as an enhancer-specific (but not promoter-specific) MYC binding site. We generated density plots of EBOX and TRE canonical sequences (CACGTG and TGA(G/C)TCA, respectively) for the K562 cell line (Fig. 2g-i). In line with established patterns of EBOX motif enrichment (Fig. 1b, 2e-f), the CACGTG sequence was prevalent in MYC’s genome-wide, promoter, and enhancer binding regions (Fig. 2g-i). Following TRE enrichment patterns in (Fig. 1b, 2e-f), the TGA(G/C)TCA sequence was prevalent in MYC’s genome-wide and enhancer binding regions, but not promoter regions (Fig. 2g-i). The central localization (Feldker et al., 2020; Lorenzin et al., 2016; Pellanda et al., 2021) of CACGTG sequence in (Fig. 2g-i) confirms that MYC binding is driven by EBOX sequence genome-wide, at promoters, and at enhancers. Similarly, the central localization of the TGA(G/C)TCA sequence in (Fig. 2i) confirms that MYC binding is driven by TRE sequence at enhancers.

### MYC occupancy of TRE site at enhancers preferentially increases with increasing MYC levels

MYC oncogene levels are often elevated in various human cancer types (Dhanasekaran et al., 2022; Lourenco et al., 2021), leading to altered transcriptional properties of MYC in cancer cells (Jakobsen & Siersbæk, 2025; Kress et al., 2016; Kress et al., 2015; Lorenzin et al., 2016; Sabò et al., 2014; Walz et al., 2014). Hence, we investigated whether MYC expression levels could also influence the extent of MYC binding to TRE sites through synergy with AP-1. We analyzed ChIP-seq datasets from the U2OS cell line with Doxycycline (Dox)-inducible MYC expression (Muthalagu et al., 2014; Walz et al., 2014) (Fig. 3a). In the absence of Dox (−Dox), U2OS cells maintain low endogenous MYC levels (low-MYC conditions) (Muthalagu et al., 2014; Walz et al., 2014). However, upon Dox addition (+Dox), they exhibit overexpressed MYC levels (high-MYC conditions) (Muthalagu et al., 2014; Walz et al., 2014). The number of peaks in MYC ChIP-seq of U2OS cells increased with the addition of Dox (Fig. 3b), transitioning from low-MYC to high-MYC conditions.

**Fig. 3:**
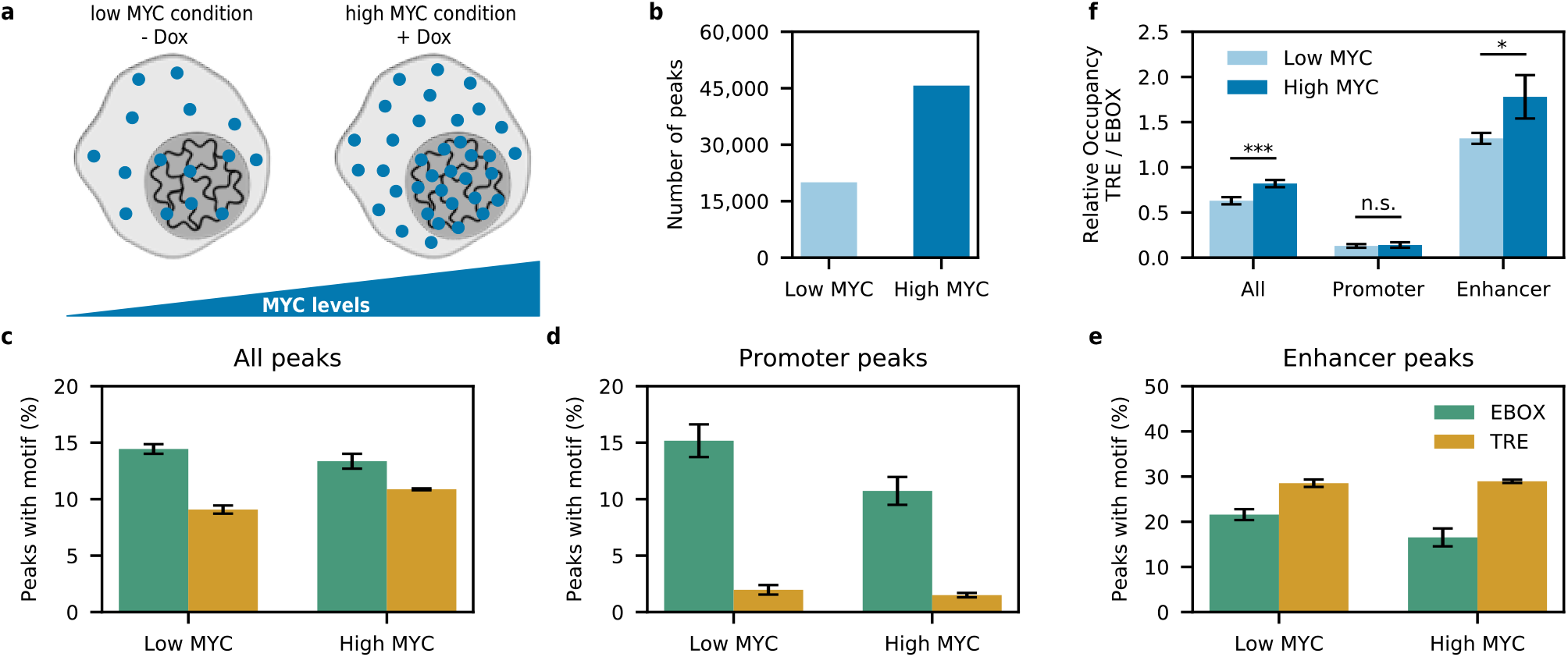
Using U2OS MYC ChIP-seq datasets to show increase in MYC binding to EBOX and TRE sites with increase in MYC levels. **a.** Schematic depicting Doxycycline inducible MYC expression in the U2OS cell line. **b**. Number of MYC ChIP-seq peaks identified from MYC ChIP-seq datasets of U2OS cell line under low- (−Dox) and high-(+Dox) MYC conditions. **c-e**. Enrichment of EBOX and TRE motifs in **c**. all genomic regions, **d**. promoters, and **e**. enhancers bound by MYC in the U2OS cell line as a function of MYC levels. **f**. Variation in relative occupancy (TRE/EBOX) as a function of MYC levels in all genomic regions, promoters and enhancers bound by MYC in the U2OS cell line. P-value calculated using two-sided Welch’s t-test. ***p<0.001, *p<0.05, and n.s is not significant. **c-f**. Data represents mean and standard deviation of 4 replicates.

When we performed *de novo* motif discovery on these low- and high-MYC ChIP-seq datasets, we found that the enrichment trends of EBOX and TRE motifs were consistent with previously established patterns of MYC binding observed in MYC ChIP-seq analysis from the ENCODE database (Fig. 1, 2) (Dunham et al., 2012; Hitz et al., 2023; Kagda et al., 2023; Luo et al., 2019). Specifically, both EBOX and TRE motifs were significantly enriched in low- and high-MYC conditions (Fig. 3c). Furthermore, the EBOX motif was significantly enriched in both MYC-bound promoters and enhancers, whereas the TRE motif was highly abundant exclusively in MYC-bound enhancers but not in promoters (Fig. 3d,e). These enrichment patterns were observed under both low- and high-MYC conditions.

To study how MYC binding patterns change with varying MYC expression levels, we calculated the occurrence of *de novo* enriched EBOX and TRE motifs in low- and high-MYC ChIP-seq datasets. There was an increase in the occurrence of EBOX and TRE motifs with increasing MYC levels (Extended Data Fig. 3a-c). This trend was consistent across MYC’s genome-wide, promoter, and enhancer binding regions. Although the occurrence of TRE and EBOX motifs increased with increase in MYC expression levels, TRE still remains an enhancer-specific MYC binding site independent of MYC levels in cells (Fig. 3e).

While the general motif enrichment trends were consistent across varying MYC expression levels, we noticed differences in the relative enrichment of TRE motifs compared to EBOX motifs between low and high MYC levels (Fig. 3c,d,e). To quantify this difference, we defined a metric called relative occupancy (RO), representing the ratio of TRE motif enrichment to EBOX motif enrichment (Fig. 3f). We examined the RO values across all, promoter, and enhancer MYC binding regions. For all MYC-bound genomic regions, we observed that the RO value was less than one under both low- and high-MYC conditions. This finding implies that, overall, direct binding of MYC to DNA through EBOX sites occurs more frequently than MYC binding to TRE sites through synergy with AP-1. At promoters, the RO value was considerably less than one under both low- and high-MYC conditions. Thus, MYC primarily utilizes promoters to directly bind to DNA through EBOX sites rather than binding to TRE sites through synergy with AP-1. In contrast, at enhancers, the RO value was greater than one under both low- and high-MYC conditions. This suggests that, at enhancers, MYC binding to TRE sites through synergy with AP-1 occurs more frequently than direct binding of MYC to DNA through EBOX sites.

Moreover, RO values significantly increased with increasing MYC levels for all MYC-bound genomic regions and MYC-bound enhancers. However, at MYC-bound promoters, RO values did not significantly change with MYC levels. This finding indicates that, as MYC levels increase, MYC preferentially binds to TRE sites through synergy with AP-1 rather than directly binding to EBOX sites. Notably, this increase in MYC binding to TRE sites is mostly localized to enhancers.

### MYC utilizes the TRE binding site at enhancers to regulate specific biological processes

Previous analyses revealed that MYC occupies TRE sites specifically at enhancers. However, the biological significance of MYC binding to TRE at enhancers still remains to be determined. To help address this question, we performed two types of analyses. First, to gain a comprehensive understanding of MYC’s transcriptional network, we studied all genes directly bound by MYC in low- and high-MYC ChIP-seq datasets (Fig. 4a). Genes were classified as MYC “targets” only if they contained a MYC ChIP-seq peak at their promoters. MYC target genes identified through the analysis correspond to potential candidate genes that can be regulated upon MYC binding.

**Fig. 4:**
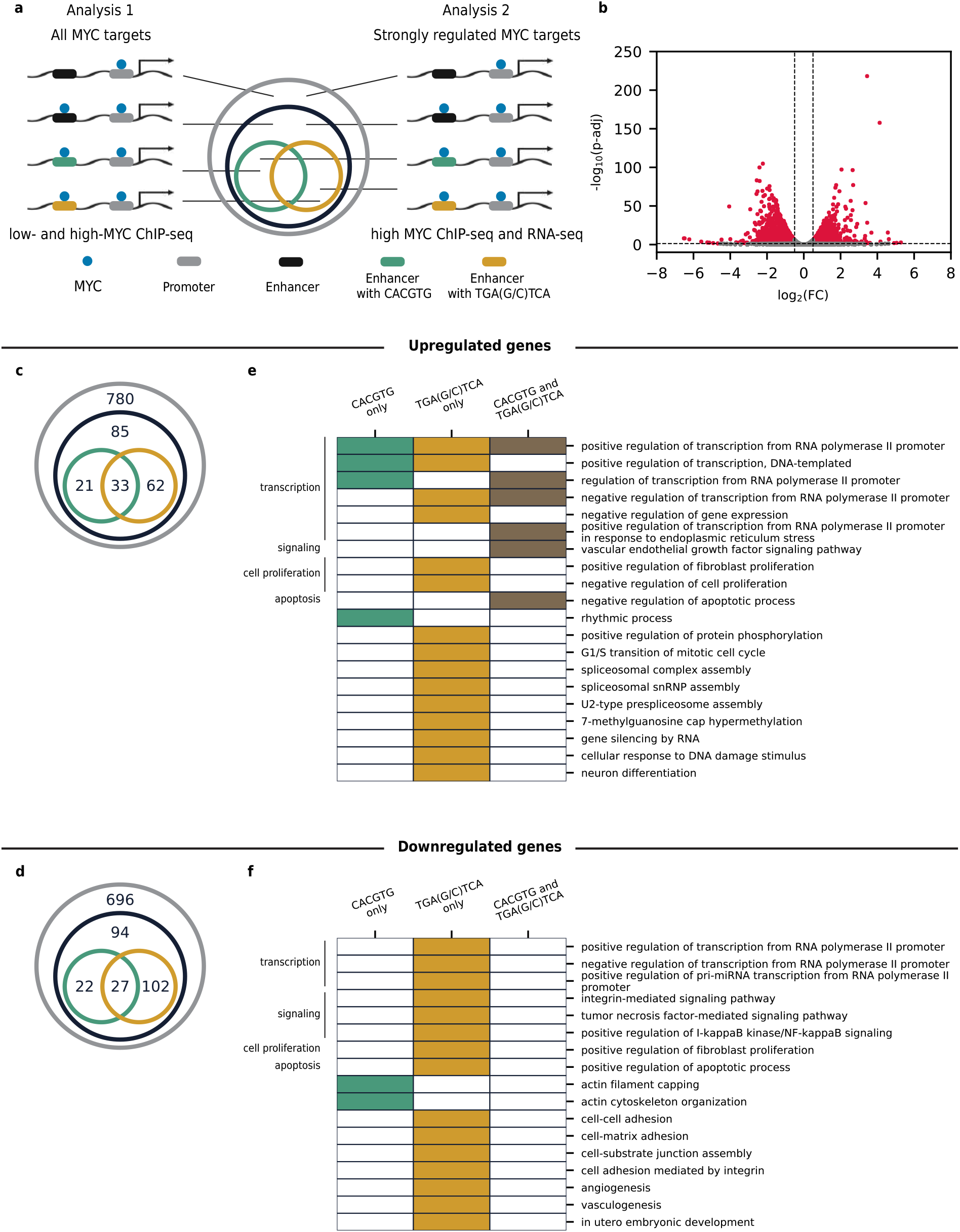
Identifying distinct biological processes regulated by MYC binding to TRE sites at enhancers by performing Gene Ontology. **a.** Schematic depicting classification of MYC target genes into subsets based on the enhancer binding patterns of MYC in ChIP-seq of the U2OS cell line. The outermost grey circle in the venn diagram represents MYC target genes set, i.e. genes containing MYC ChIP-seq peak at their promoters. Black circle represents a subset of MYC target genes which contain an additional MYC ChIP-seq peak at their enhancers. Green (or yellow) circle represents MYC target gene subsets, specifically containing a CACGTG (or TGA(G/C)TCA) sequence within their enhancer MYC ChIP-seq peak. In Analysis 1, MYC target genes were identified based on MYC binding to genes in low- and high-MYC ChIP-seq datasets of the U2OS cell line. In Analysis 2, strongly regulated MYC target genes were identified based on MYC binding to genes in high-MYC ChIP-seq, and differential expression of those genes in +/-Dox RNA-seq of the U2OS cell line. **b-f:** Results for Analysis 2. **b**. Volcano plot of differentially expressed genes identified using RNA-seq in U2OS cell line. Genes that are significantly differentially expressed (−0.5<log2(FC)<0.5 and p-adj<0.05) are highlighted in red. Fold change (FC) was calculated as the ratio of gene expression in +Dox condition over gene expression in -Dox condition. **c-d**. Number of genes bound by MYC in high-MYC condition, and genes significantly **c**. up- or **d**. downregulated upon Dox addition identified using ChIP-seq and RNA-seq data of U2OS cell line. **e-f:** Gene Ontology analysis of genes in green and yellow subsets identified in panels **c** and **d** respectively.

**Fig. 5:**
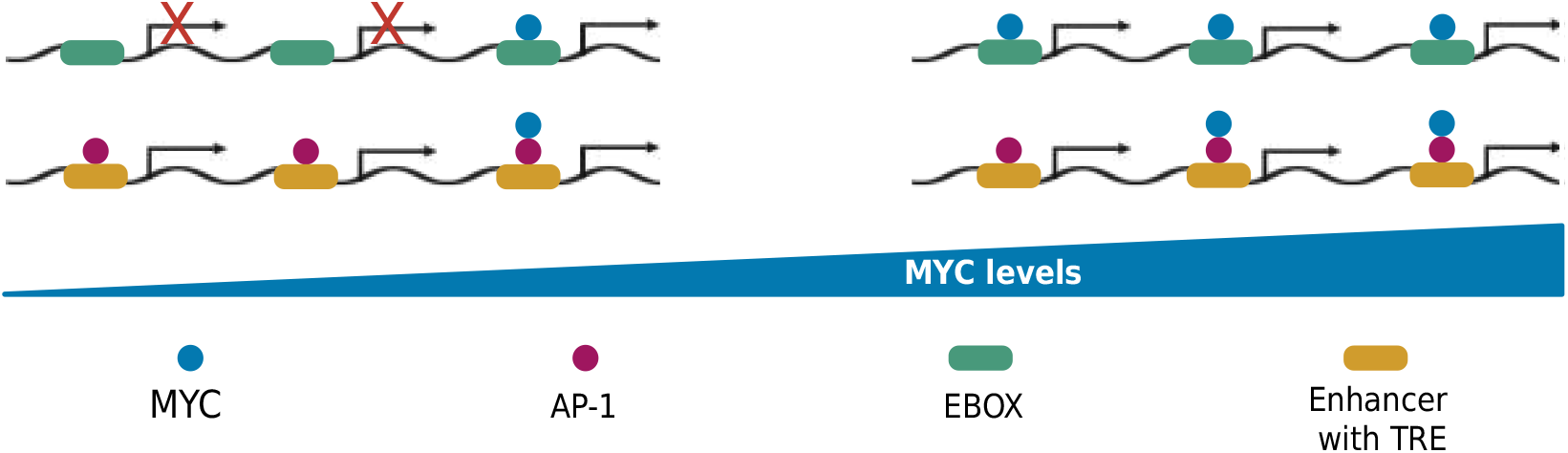
Schematic showing study findings where MYC DNA-binding patterns and MYC synergy with AP-1 alter as a function of MYC expression levels in cells. These mechanisms contribute to transcriptional rewiring and ultimately cellular reprogramming in MYC-driven cancerous cells.

The identified MYC target genes were then divided into five gene sets (Fig. 4a) (see Supplemental Methods). The broadest set corresponds to all MYC target genes (grey). Within this set, genes that contained an additional MYC ChIP-seq peak at their enhancers were selected (black). These genes were further analyzed to identify exact matches for EBOX (CACGTG) (green) or TRE (TGA(G/C)TCA) (yellow) sequences within their enhancer MYC ChIP-seq peaks. Target genes with MYC-bound TRE at enhancers (yellow) and those with MYC-bound EBOX at enhancers (green) were selected for further analysis to characterize the role played by MYC binding to TRE and EBOX at enhancers. These two gene sets correspond to three types of MYC enhancer elements (EEs): EBOX-only, TRE-only, and both EBOX and TRE (referred to as EBOX-TRE). Gene Ontology (GO) analysis (Huang da et al., 2009; Sherman et al., 2022) was performed separately for genes associated with each of the EEs to identify their associated biological processes (Extended Data Fig. 4c,d).

We observed that when MYC levels go from low to high, the number of genes bound to each subset increased (Extended Data Fig. 4a,b); however, comparing the GO tables between low- (Extended Data Fig. 4c) and high-MYC conditions (Extended Data Fig. 4d) revealed distinct binding patterns. Under low-MYC conditions, MYC regulated genes by binding to EBOX-only and TRE-only EEs (Extended Data Fig. 4c). In contrast, under high-MYC conditions, MYC also regulated genes by binding to EBOX-TRE EE, indicating the additional cellular functions mediated by MYC-AP1 interactions (Extended Data Fig. 4d).

Several processes were regulated in both low- and high-MYC conditions but exhibited different binding patterns (Extended Data Fig. 4c,d). In low-MYC conditions (Extended Data Fig. 4c), transcription-related processes were regulated by MYC binding to TRE-only EE, whereas in high-MYC conditions (Extended Data Fig. 4d), these processes were also regulated by MYC binding to EBOX-only and EBOX-TRE EEs. Phosphorylation and signaling processes were regulated by MYC binding to EBOX-only and TRE-only EEs under low-MYC conditions (Extended Data Fig. 4c); however, in high-MYC conditions (Extended Data Fig. 4d), these processes were regulated by MYC binding to TRE-only and EBOX-TRE EEs. Cell cycle processes gained EBOX-only EE in addition to TRE-only EE when MYC levels increased from low to high (Extended Data Fig. 4c,d). However, cell proliferation, apoptosis, and cell migration remained regulated by TRE-only EE in both low- and high-MYC conditions (Extended Data Fig. 4c,d).

Distinct biological processes were also regulated by MYC binding to various EEs across low- and high-MYC conditions (Extended Data Fig. 4c,d). For instance, in low-MYC condition (Extended Data Fig. 4c), spliceosomal snRNP assembly and positive cyclin-dependent kinase activity were regulated by MYC binding to TRE-only EE. In contrast, under high-MYC conditions (Extended Data Fig. 4d), embryonic development was regulated by TRE-only EE. Additionally, under high-MYC conditions (Extended Data Fig. 4d), angiogenesis, receptor internalization, and Schwann cell development were regulated by MYC binding to EBOX-TRE EE.

In order to study genes strongly regulated by MYC in cancerous conditions, we next analyzed published RNA-seq data to identify genes that show significant differential expression upon MYC induction, focusing specifically on genes directly bound by MYC in the high-MYC ChIP-seq dataset (Fig. 4a). The strongly up- and downregulated MYC target genes identified were then classified into different subsets as previously described (Fig. 4a). GO analysis was performed on these target gene subsets associated with different EEs to identify the biological processes modulated by them (Fig. 4e,f) (see Supplemental Methods).

The number of genes bound to each subset is comparable between upregulated and downregulated genes, except for TRE-only EE, which MYC uses to downregulate more genes than to upregulate (Fig. 4c,d). In both upregulated and downregulated cases (Fig. 4e,f), transcription, signaling, cell proliferation, and apoptosis were identified as commonly regulated processes. A comparison of enhancer binding patterns for these processes between upregulated and downregulated cases (Fig. 4e,f) revealed that downregulated GO terms primarily contained TRE EE. In contrast, upregulated GO terms (Fig. 4e) contained all three EEs and exhibited distinct binding patterns. Interestingly, we observed that endoplasmic reticulum (ER) stress related transcription was upregulated by MYC binding to EBOX-TRE EE (Fig. 4e), which may potentially represent a mechanism of stress response regulation by MYC (Tameire et al., 2019).

The up- and downregulated cases also contained various distinct GO-enriched terms (Fig. 4e,f). Rhythmic process-related genes were upregulated through MYC binding to EBOX-only EE (Fig. 4e). Several other processes, including protein phosphorylation, cell cycle, DNA damage response, and neuron differentiation were upregulated by MYC binding to TRE-only EE (Fig. 4e). Interestingly, actin-related processes were found to be downregulated through MYC binding to EBOX-only EE (Fig. 4f). Additionally, several more GO terms, including cell adhesion, angiogenesis, and embryonic development were downregulated through MYC binding to TRE-only EE (Fig. 4f).

## Discussion

In this work, we leverage publicly available next-generation sequencing (NGS) datasets (Dunham et al., 2012; Hitz et al., 2023; Kagda et al., 2023; Luo et al., 2019; Muthalagu et al., 2014; Walz et al., 2014) to uncover a novel mechanism that MYC may utilize to drive and sustain various types of human cancers. Using ChIP-seq, we demonstrate that TRE serves as an indirect enhancer binding site for MYC, enabling its synergistic interaction with AP-1 TFs and the modulation of its target gene expression. This synergistic interaction between major oncogenes, MYC and AP-1 families (Eferl & Wagner, 2003; Garces de los Fayos Alonso et al., 2018; Jakobsen & Siersbæk, 2025; Kress et al., 2015), is relevant for both MYC-driven as well as AP-1 driven cancers. Specifically, we study the broad DNA binding patterns of MYC by analyzing published MYC ChIP-seq datasets. Through *de novo* motif discovery (Heinz et al., 2010) on ENCODE MYC ChIP-seq datasets (Dunham et al., 2012; Hitz et al., 2023; Kagda et al., 2023; Luo et al., 2019), we reveal the co-enrichment of the AP-1 family binding motif (TRE) (Eferl & Wagner, 2003; Garces de los Fayos Alonso et al., 2018), in addition to the enrichment of the known MYC consensus binding motif (EBOX) (Jakobsen & Siersbæk, 2025; Kress et al., 2015) (Fig. 1a,b) in 8 out of 11 analyzed cell lines. Previously, different studies have reported the enrichment of the TRE motif in MYC ChIP-seq datasets (Lu et al., 2016; Wang et al., 2012), the co-association between MYC and AP-1 genomic binding regions (Dunham et al., 2012; Guo et al., 2014; Roca et al., 2014), as well as protein-level interactions between MYC and the AP-1 TF JUN using co-immunoprecipitation (Levy & Forman, 2010). However, the connection between these findings, and their biological significance has not been investigated. Through analyses integrating MYC and various AP-1 family TF ChIP-seq datasets (Dunham et al., 2012; Hitz et al., 2023; Kagda et al., 2023; Luo et al., 2019) across 7 cell lines, we demonstrate that the co-enriched TRE motif in MYC ChIP-seq datasets represents an indirect DNA binding site for MYC, occupied in synergy with AP-1 TFs (Fig. 1c-f).

Our report reveals a novel synergy between MYC and AP-1 TFs in DNA binding through TRE sites across multiple cell lines. Our *de novo* motif enrichment analysis across 7 cell lines reveals that the TRE motif is preferentially enriched at enhancers rather than promoters (Fig. 2e,f), suggesting that the TRE binding site plays an enhancer-specific role in MYC function. This finding indicates that MYC, in synergy with AP-1, may utilize the TRE sites at enhancers to modulate gene expression from basal levels in various cancer cell lines. Through *de novo* motif discovery on MYC ChIP-seq datasets at both low- and high-MYC expression levels (Muthalagu et al., 2014; Walz et al., 2014) (Fig. 3a,b), we observe an increased extent of MYC binding to TRE compared to EBOX at high-MYC conditions relative to low-MYC conditions (Fig. 3f). This finding suggests that at high MYC levels, TRE sites at enhancers could become a key mechanism for MYC-mediated gene regulation in cancers.

Our genome-wide analysis reveals that TRE motif enrichment in MYC ChIP-seq occurs in a cell-type specific manner, observed in 8 out of 11 analyzed cell lines (Fig. 1a-b). This cell-type specificity aligns with our finding that MYC-AP-1 interactions occur at TRE sites with enhancer functions, as enhancers are known to operate in a cell-type specific manner (Heinz et al., 2015; Schoenfelder & Fraser, 2019). However, both MYC and AP-1 TFs are ubiquitously expressed (Köster et al., 2023; Liu et al., 2006; Llombart & Mansour, 2022) across most cell types, albeit to varying degrees. Despite the cell-type specificity of MYC-AP-1 synergy identified in this work, the widespread expression of these TFs (Köster et al., 2023; Liu et al., 2006; Llombart & Mansour, 2022) suggests that this interaction could emerge when MYC and AP-1 expression levels are altered, through various genomic (Dhanasekaran et al., 2022; Fittall et al., 2018; Meyer & Penn, 2008; PANAGOPOULOS & HEIM, 2022), epigenetic (Fatma et al., 2022; Joo et al., 2016; Schuijers et al., 2018), and even microenvironment (Benaud & Dickson, 2001; Chiariello et al., 2001; Cowles et al., 2000; Peake et al., 2000; Shynlova et al., 2002) potentiated changes during cancer initiation and progression (Hanahan & Weinberg, 2000; Hanahan & Weinberg, 2011). Hence, MYC-AP-1 synergy has the potential to be co-opted in diverse cell types and cancers, extending MYC’s well appreciated role of a cell-intrinsic driver (Dhanasekaran et al., 2022; Gabay et al., 2014).

By integrating ChIP-seq and RNA-seq datasets of differential MYC expression (Muthalagu et al., 2014; Walz et al., 2014) (Fig. 4a,b), we identify several target genes regulated by MYC binding to enhancers with TRE sites (Fig. 4c-d). Our results reveal that when MYC levels increase, more of these target genes are downregulated than upregulated (Fig. 4c,d), suggesting that MYC binding to TRE enhancer sites may play a more specific role in gene downregulation than upregulation. Additionally, we demonstrate that MYC utilizes TRE binding sites at enhancers to regulate a multitude of cellular processes including transcription, signaling, proliferation, apoptosis, cell cycle, phosphorylation, and cell adhesion (Fig. 4e,f). Notably, several of these identified processes correspond to well-known cancer hallmarks (Hanahan & Weinberg, 2000; Hanahan & Weinberg, 2011), indicating that gene regulation via MYC binding to TRE sites at enhancers plays a key role in driving and sustaining cancers.

Although TRE is a well-established binding motif for the AP-1 TF family in mammalian cell lines (Eferl & Wagner, 2003; Garces de los Fayos Alonso et al., 2018), its role in MYC-mediated gene regulation has not been identified previously. This is likely because TRE functions as an enhancer binding site for MYC (Fig. 2e-f), which is often located at a considerable distance from its target genes (Heinz et al., 2015; Schoenfelder & Fraser, 2019), making its identification inherently challenging (Heinz et al., 2015; Schoenfelder & Fraser, 2019). Our work reveals a novel finding that MYC synergistically interacts with multiple (at least 4) AP-1 TF family members to occupy DNA at TRE binding sites within enhancers (Fig. 1c-f, 2). The AP-1 TF family consists of many members that dimerize with each other and bind to TRE sites (Eferl & Wagner, 2003; Garces de los Fayos Alonso et al., 2018). Further work is needed to determine the contribution of individual AP-1 family members and their respective dimers to this synergy with MYC. A previous study provided evidence of MYC forming a complex with the AP-1 TF member JUN through co-immunoprecipitation (co-IP) (Levy & Forman, 2010). Co-IP between all AP-1 TFs and MYC can help identify individual AP-1 family members that form complexes with MYC in cells. Subsequent studies can then determine whether the interaction between MYC and AP-1 observed in this work is part of a regulatory complex or a direct protein-protein interaction.

## Acknowledgements

This work was supported in part by NIH grants CA250044, EB01775309 and the Center for Precision Engineering for Health (CPE4H) at the University of Pennsylvania. We acknowledge the ENCODE Consortium and the ENCODE production laboratory(ies) generating the particular dataset(s) utilized in this work. Certain figure panels (3a, 4a, and 5) in this work were generated using BioRender (https://BioRender.com). Computational resources were provided in part by the School of Engineering and Applied Sciences at the University of Pennsylvania and the Pittsburgh Supercomputing Center through Access allocation MCB200101. This work also used the Galaxy web platform (https://usegalaxy.org/) for analysis. We thank members of Lim and Radhakrishnan labs for helpful discussions.

## Author contributions

Data analysis: R.K.; Conceptualization: R.K., R.R., and B.L.; Methodology: R.K., R.R., and B.L.; Investigation: R.K., R.R., and B.L.; Writing – original draft: R.K., R.R., and B.L.; Writing – review and editing: R.K., R.R., and B.L.; Supervision: R.R., and B.L.; Project administration: R.R., and B.L.; Funding acquisition: R.R., and B.L.

## Extended Data figure legends

**Extended Data Fig. 1: a**. Overlap of MYC and various AP-1 TF DNA binding regions identified using ChIP-seq datasets for multiple cell lines. P-value of overlap calculated using a hypergeometric test. **b-c**. Occurrence of **b**. EBOX and **c**. TRE motifs before and after the removal of co-bound peaks with various AP-1 family TFs in multiple cell lines. The drastic reduction in MYC ChIP-seq peaks containing TRE motifs upon removal of peaks overlapping with AP-1 ChIP-seq shows TRE motifs enriched in MYC ChIP-seq as a synergistic binding site of MYC with AP-1 TFs.

**Extended Data Fig. 2: a-b**. Heatmaps showing ChIP-seq binding intensities around promoter and enhancer regions of MYC for various cell lines analyzed. **a**. For promoters, MYC binding intensities are plotted around the center of TSS of actively transcribed genes bound by MYC. Rows are sorted in decreasing order of MYC binding intensities. **b**. For enhancers, MYC and H3K27ac binding intensities are plotted around the center of active enhancers bound by MYC. Rows are sorted in decreasing order of H3K27ac binding intensities. All heatmaps cover +/-5 kb window around the reference chosen for centering. Color bar indicates ChIP-seq binding intensities for corresponding heatmaps.

**Extended Data Fig. 3: a-c**. Occurrence of EBOX and TRE motifs in **a**. all genomic regions, **b**. promoters, and **c**. enhancers bound by MYC in the U2OS cell line as a function of MYC levels. Data represents mean and standard deviation of 4 replicates.

**Extended Data Fig. 4: Identifying distinct biological processes regulated by MYC binding to TRE sites at enhancers by performing Gene Ontology. a-d:** Results for Analysis 1. **a-b**. Number of genes bound by MYC in **a**. low- and **b**. high-MYC condition identified using ChIP-seq data of U2OS cell line. **c-d:** Gene Ontology analysis of genes in green and yellow subsets identified in panels **a** and **b** respectively.

