## Supplemental Methods for "MYC and AP-1 oncogenes synergistically bind enhancers to rewire transcription"

**List of tools:**

1. HOMER software suite (v 4.11.1) (Heinz et al., 2010)
2. Bedtools (v 2.31.0) (Quinlan & Hall, 2010)
3. IGV genome browser (web application) (v 2.2.7) (Robinson et al., 2011; Thorvaldsdóttir et al., 2012)
4. IntAct Molecular Interaction Database Portal (v 1.0.4) (del Toro et al., 2021)
5. DAVID functional annotation tool (v 2023q4) (Huang da et al., 2009; Sherman et al., 2022)
6. deepTools (v 3.5.3) (Ramírez et al., 2016)
7. bioRender
8. Python (v 2.7.16)
9. Adobe Illustrator

**Motif discovery:** *de novo* motif discovery was performed on ChIP-seq datasets (in BED format) using the findMotifsGenome.pl program from the HOMER software suite. Reference genomes (hg19, hg38, or mm10) were specified according to the databases from which the ChIP-seq files were obtained. Enriched motifs were identified within a +/-100 bp region around the center of ChIP-seq peaks. The number of background sequences was set to be 5 times the total number of ChIP-seq peaks. The HOMER software suite generated the random background sequences as controls for the motif discovery runs, as described in HOMER documentation. The motif length was set to 10 bp. The enrichment value of a motif was calculated as the percentage enrichment in target sequences minus the percentage enrichment in background sequences. Each motif discovery run was performed 4 times, and the mean and standard deviation of enrichment values are reported in the figures. The number of peaks containing an enriched motif (referred to as “occurrence”) was calculated by multiplying the mean enrichment value (in %) by the total number of ChIP-seq peaks. Statistical significance of the *de novo* enriched motif was calculated by the HOMER software using a binomial test. Only significantly enriched motifs (p-value < 1e-60) were used for analysis.

**ChIP-seq overlap:** The mergePeaks.pl program in the HOMER software suite was used to identify overlapping peaks between two ChIP-seq datasets (in BED format). Peaks were considered overlapping if the distance between their centers was less than 200 bp. The genome size was set to the total length (in bp) of the reference genome of the ChIP-seq datasets. Statistical significance of overlap between ChIP-seq datasets was calculated by the mergePeaks program using a hypergeometric test.

**ChIP-seq subtraction analysis:** Requirement of MYC-AP-1 synergy for MYC binding to the TRE motif was demonstrated by comparing the change in occurrence of TRE (and EBOX) motifs in all MYC ChIP-seq peaks versus MYC ChIP-seq peaks that do not overlap with individual AP-1 transcription factor ChIP-seq peaks. Overlapping peaks between MYC and various AP-1 family transcription factors were removed from MYC ChIP-seq datasets (in BED format) using the bedtools subtract utility with the -A option. *de novo* motif discovery was then performed on all MYC ChIP-seq peaks (or MYC ChIP-seq peaks that do not overlap with AP-1 ChIP-seq peaks) to determine the occurrence of enriched EBOX and TRE motifs as described previously. The mean occurrences calculated from 4 replicates of motif discovery runs were used to generate the figure panels. Within a cell line, motif occurrences in MYC ChIP-seq peaks that do not overlap with AP-1 ChIP-seq peaks were normalized to motif occurrences in all MYC ChIP-seq peaks, and the results were presented as a heatmap.

**IGV plotting:** IGV tracks were generated by loading ChIP-seq files (in bigWig format) into the IGV genome browser. Reference genomes (hg19, hg38, or mm10) were specified according to the databases from which the ChIP-seq files were obtained. Binding profiles of MYC, H3K4me3, or H3K27ac were visualized by zooming into specific genomic regions.

**Promoter Enhancer separation:** Promoter identification (Lin et al., 2012): Transcribed genes were defined as those with an H3K4me3 ChIP-seq peak within +/- 5 kb of the TSS. MYC-bound promoters were identified as MYC ChIP-seq peaks within +/- 1 kb of the TSS of transcribed genes. Enhancer identification (Lin et al., 2012): Active enhancer regions were defined as H3K27ac ChIP-seq peaks located outside promoter regions (more than 5 kb away from the TSS). MYC-bound enhancers were identified as MYC ChIP-seq peaks within +/- 1 kb of the center of active enhancer regions.

For the identification of promoter and enhancer regions of MYC binding, the following tools were utilized. The annotatePeaks.pl program in the HOMER software suite was used to obtain the nearest gene corresponding to a ChIP-seq peak, and the distance between the ChIP-seq peak and the TSS of the nearest gene. Information regarding the TSS of various genes was obtained from the genome.tss file included in the HOMER software suite. The bedtools subtract utility with -A option was used to remove overlapping peaks between two ChIP-seq datasets. The bedtools window utility was used to select ChIP-seq peaks from one dataset that are within a specific window (in bp) of another ChIP-seq dataset.

**Promoter enhancer heatmaps:** Heatmaps for promoters (Lin et al., 2012) were generated by plotting the MYC ChIP-seq signal intensity over MYC-bound transcriptionally active promoters, with each row representing a +/- 5 kb region centered on the TSS. The rows were arranged in descending order of MYC ChIP-seq signal intensity, and color bars indicate the intensity of the MYC ChIP-seq signal. Heatmaps for enhancers (Lin et al., 2012) were generated by plotting the H3K27ac or MYC ChIP-seq signal intensity over MYC-bound active enhancer regions, with each row representing a +/- 5 kb region centered on the active enhancers. The rows were arranged in descending order of H3K27ac ChIP-seq signal intensity, and color bars indicate the intensity of the H3K27ac or MYC ChIP-seq signal. The computeMatrix tool from deepTools suite was used in reference-point mode to generate intermediate matrix files corresponding to each heatmap. These matrix files were then visualized as heatmaps using the plotHeatmap tool from deepTools suite.

**Motif density plots:** The annotatePeaks.pl program in the HOMER software suite was used to generate density plots of EBOX (CACGTG) and TRE (TGA(G/C)TCA) sequences in the ChIP-seq datasets. CACGTG and TGA(G/C)TCA sequences were scanned within +/-300 bp around the center of the ChIP-seq peaks. The bin size of the generated histogram was set to 10 bp.

**Relative occupancy:** Relative occupancy (RO) was defined as the ratio of TRE motif enrichment to EBOX motif enrichment in the analyzed ChIP-seq dataset. RO was calculated for 4 replicates of motif discovery runs. The statistical significance of change in RO between low-MYC and high-MYC ChIP-seq datasets was calculated using two-sided Welch's unequal variances t-test.

**Gene Ontology**

Gene Ontology (GO) analysis (Huang da et al., 2009; Sherman et al., 2022) was performed on MYC-regulated genes to identify the biological significance of MYC binding to TRE sites at enhancers. MYC-regulated genes were identified using ChIP-seq and RNA-seq datasets from the U2OS cell line (Muthalagu et al., 2014; Walz et al., 2014). In this analysis, genes were classified as MYC “target genes” only if they contained a MYC ChIP-seq peak at their promoter. Here, the gene sets and biological processes corresponding to the novel MYC-enhancer binding site, TRE, are studied in comparison to the known MYC binding site, EBOX.

Two types of GO analysis were conducted in this study. For each analysis type, MYC target genes were defined as:

1. Analysis type 1: Genes bound by MYC (Fig. 4a left), identified using low- and high-MYC ChIP-seq datasets.
2. Analysis type 2: Genes bound by MYC and showing significant differential expression (Fig. 4a right), identified using high-MYC ChIP-seq and +/-Dox RNA-seq datasets.

Identification of MYC Target Genes:

For each GO analysis type, five types of MYC target gene sets were defined, as outlined below, and represented using a Venn diagram schematic (Fig. 4a). A standard Venn diagram format was used to display the different target gene sets and subsets, where target gene sets are depicted as circles, and subsets within these target gene sets are shown as smaller circles nested within the larger circles. The number of genes corresponding to different sets are represented on the Venn diagram.

For analysis type (1), various sets in the venn diagram (Fig. 4a left) correspond to the following MYC target genes: The grey circle represents genes with MYC ChIP-seq peak at their promoters (MYC target genes). The black circle represents genes with MYC ChIP-seq peaks at their promoters and enhancers. The green circle represents genes with MYC ChIP-seq peaks at their promoters and enhancers, specifically containing an EBOX (CACGTG) sequence in the enhancer MYC ChIP-seq peak. The yellow circle represents genes with MYC ChIP-seq peaks at their promoters and enhancers, specifically containing an TRE (TGA(G/C)TCA) sequence in the enhancer MYC ChIP-seq peak. For analysis type (2), when RNA-seq data was additionally incorporated (Fig. 4a right), only target genes showing significant differential expression (|log2(fold change)| > 0.5, p-adj <0.05) were selected from the above sets.

MYC-bound promoters and enhancers were identified using MYC ChIP-seq datasets as previously described. The annotatePeaks.pl program from the HOMER software suite was employed to: (i) identify the nearest genes associated with various MYC-bound promoter and enhancer ChIP-seq peaks and (ii) search for EBOX (CACGTG) and TRE (TGA(G/C)TCA) sequence matches within MYC-bound enhancer ChIP-seq peaks.

From the various types of MYC target gene sets identified in (Fig. 4a), the following sets are used for further analysis: target genes regulated by MYC binding to TRE at enhancers (green) and target genes regulated by MYC binding to EBOX at enhancers (yellow). These two sets correspond to MYC target genes with three types of enhancer elements (EEs): (also referred to as "gene lists" from now on): i) EBOX-only, ii) TRE-only, and iii) both EBOX and TRE (referred to as EBOX-TRE).

Therefore, the following gene lists are used for further analysis.

1. Analysis type 1:
    a. Low-MYC ChIP-seq x three types of EEs
    b. High-MYC ChIP-seq x three types of EEs
2. Analysis type 2:
    a. Upregulated genes x three types of EEs
    b. Downregulated genes x three types of EEs

<u>Identification of Proteins That Physically Interact With Proteins Coded by MYC Target Genes:</u>

High-confidence direct physical interaction protein partners of proteins coded by the MYC target genes in each list were identified using the IntAct Molecular Interaction Database. The IntAct website was used with the following options to identify the interactor proteins:

1. Interactor species: Selected as homo sapiens (U2OS cell line), with only intra species interactions included.
2. Interactor types: Set to "protein" and "direct" to identify only direct protein-protein interactions.
3. MI score thresholds: Defined as (0.7 < MI score < 1) for genes in analysis type (1) and (0.5 < MI score < 1) for genes in analysis type (2).

The proteins corresponding to the identified protein-protein interactions were then downloaded from the IntAct website. Genes corresponding to these protein interactions were then added to the gene lists to identify biological pathways mediated by MYC target genes, similar to previous studies (Gitter & Bar-Joseph, 2013; Ma et al., 2006; Schaefer et al., 2013) which integrated gene expression datasets and protein-protein interaction networks to identify pathways regulated by differentially expressed genes.

Functional enrichment analysis:

Gene Ontology (GO) was then performed on the gene lists from the previous step using the DAVID functional annotation tool to identify the biological processes mediated by these genes. GO terms were considered significantly enriched if the Benjamini value was less than 0.01 for analysis type (1) and 0.05 for analysis type (2). GO terms were included for further analysis only if at least two genes corresponded to the enriched GO term.

We found that the GO enrichment results were robust to changes in the confidence (MI score) of protein-protein interactions. Specifically, the number of enriched GO terms corresponding to MYC target genes in TRE-only EEs was consistently greater than those in EBOX-only EEs, regardless of the MI score threshold set.

The enriched GO terms were then used to construct GO tables. For analysis type (1), two GO tables were generated for the low- and high-MYC conditions. For analysis type (2), two GO tables were generated for the up- and downregulated genes. Each table included three columns corresponding to the three EE types in MYC target genes: i) EBOX-only, ii) TRE-only, and iii) EBOX-TRE. Columns were highlighted if the enriched GO term was present for a specific EE type.

Gene Ontology table ordering protocol:

The following protocol was used to organize and present enriched GO terms in table format.

1. Within a GO table, GO terms corresponding to similar processes were grouped and labeled. Labeled groups include Transcription (T), Signaling (S), Phosphorylation (P), Apoptosis (A), Cell cycle (CC), Cell proliferation (CP), and Cell migration (CM).
2. Within these labeled groups, GO terms were reordered based on the number of EEs they corresponded to.
3. From these labeled groups, common terms across different conditions being compared (low- and high-MYC condition, or Up- and Down-regulated genes) were identified.
4. Common groups so identified were listed at the top of the tables, followed by distinct GO terms across the compared conditions.
5. Within the distinct GO terms, labeled groups were listed on top, and individual ungrouped GO terms were listed in the bottom.
6. The GO tables so obtained contain the complete list of enriched GO terms.
7. To generate GO tables provided in the figures, a maximum of 15 GO terms with least Benjamini values were used. (Fig. 4,e,f and Extended data Fig. 4c,d)
