## Supplemental Figure 1 for "MYC and AP-1 oncogenes synergistically bind enhancers to rewire transcription"

**a**

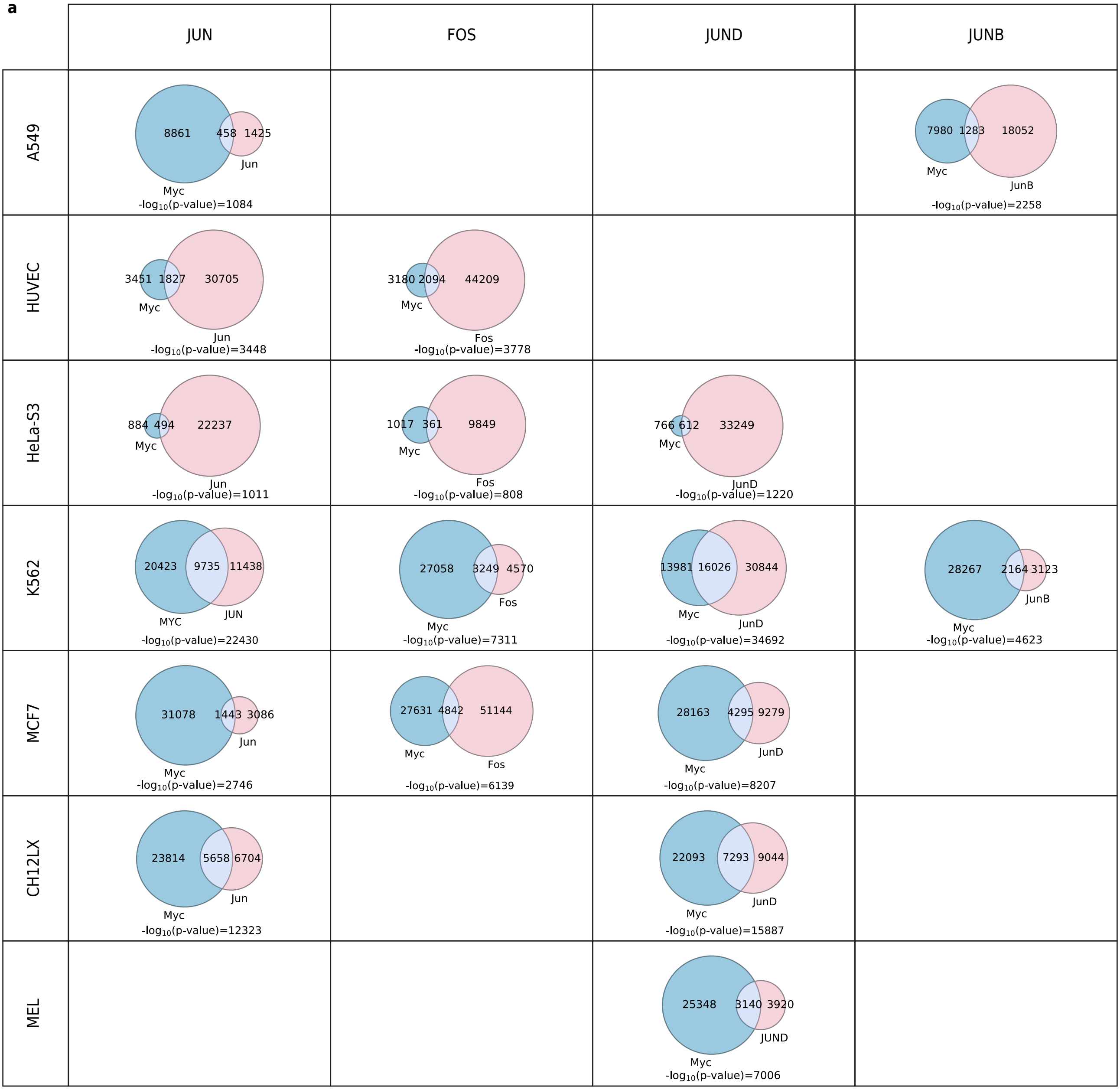


**b**

| | MYC | MYC-JUN | MYC-FOS | MYC-JUND | MYC-JUNB |
|---|---|---|---|---|---|
| A549 | 2317 | 2173 | | | 1873 |
| HUVEC | 1690 | 1131 | 1042 | | |
| HeLa-S3 | 643 | 402 | 458 | 370 | |
| K562 | 6182 | 4142 | 5436 | 3072 | 5337 |
| MCF7 | 5110 | 4512 | 4630 | 4393 | |
| CH12.LX | 6679 | 5670 | | 5405 | |
| MEL | 6411 | | | 5484 | |

**c**

| | MYC | MYC-JUN | MYC-FOS | MYC-JUND | MYC-JUNB |
|---|---|---|---|---|---|
| A549 | 841 | 656 | | | 200 |
| HUVEC | 971 | 0 | 0 | | |
| HeLa-S3 | 296 | 11 | 96 | 0 | |
| K562 | 1822 | 193 | 760 | 0 | 1041 |
| MCF7 | 1632 | 1037 | 0 | 80 | |
| CH12.LX | 1620 | 0 | | 0 | |
| MEL | 1150 | | | 225 | |
