## Supplementary figures and images for "MYC and AP-1 oncogenes synergistically bind enhancers to rewire transcription"

### Supplemental Figure 2

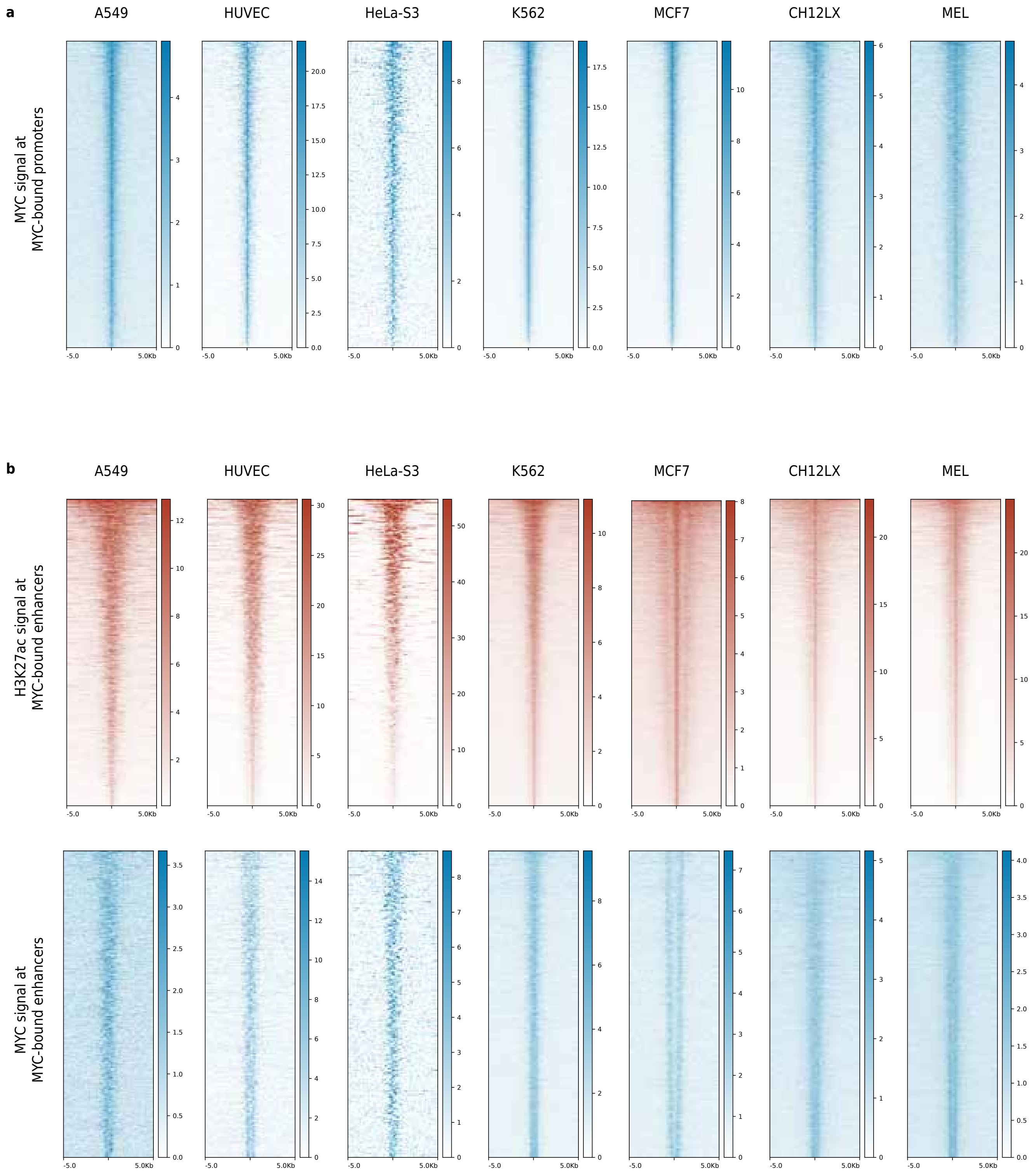

### Supplemental Figure 3

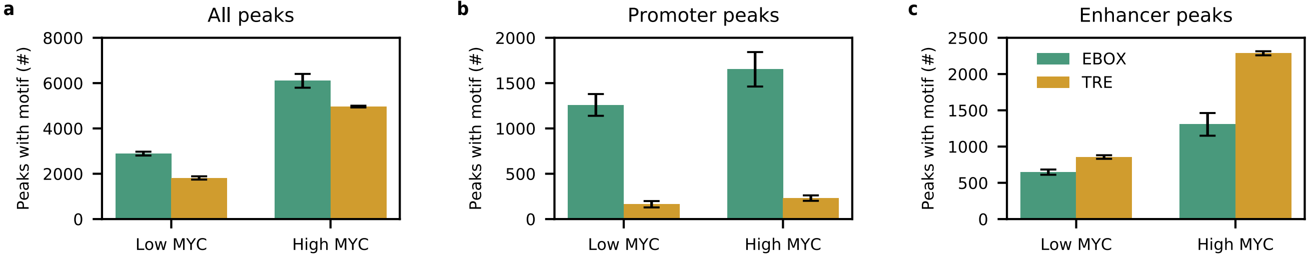

### Supplemental Figure 4

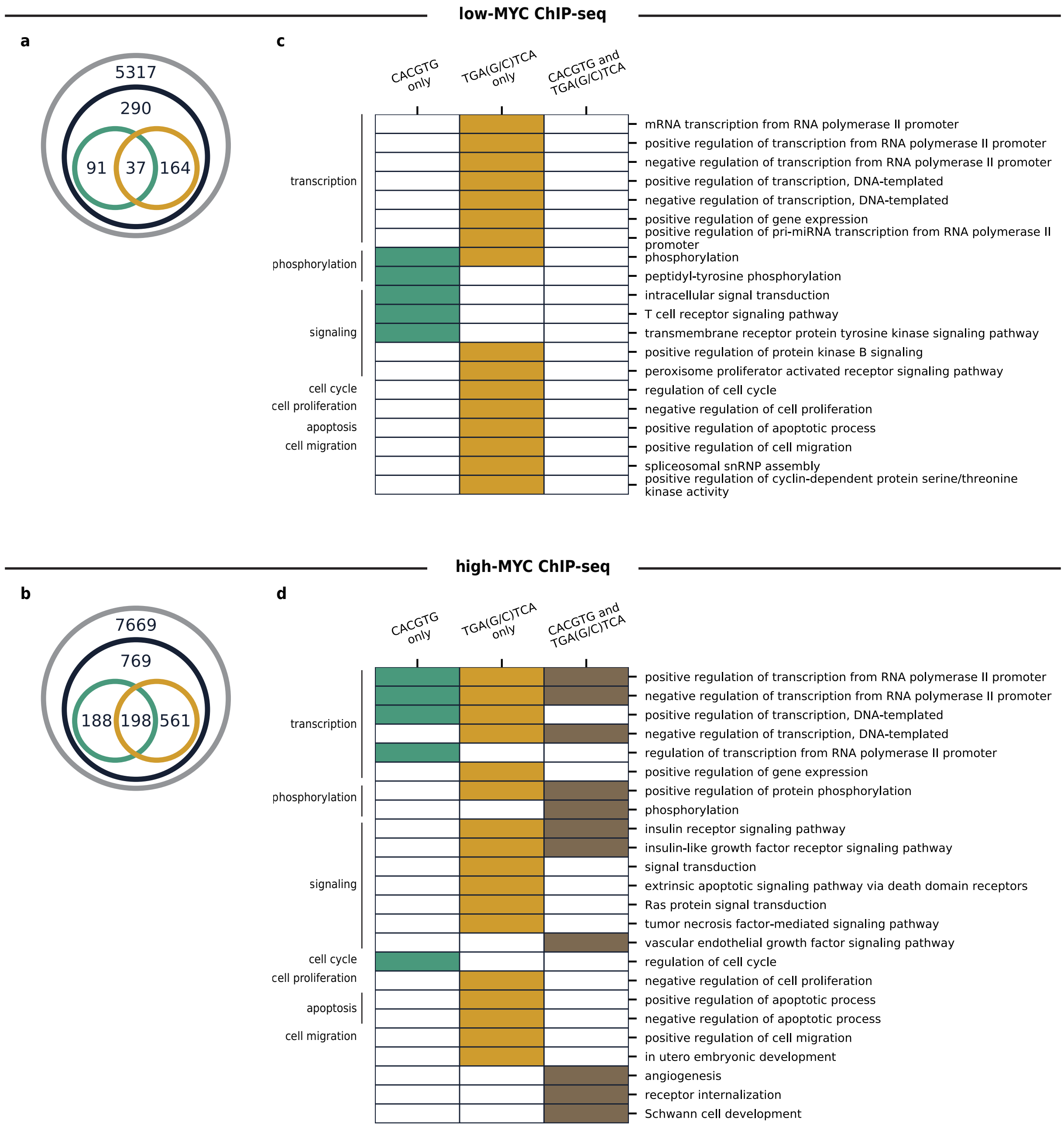
